# *Clostridium novyi* type B causes fatal hemorrhagic disease in wild Asian elephants

**DOI:** 10.1101/2024.12.31.630866

**Authors:** Soumesh Kumar Padhi, Avishek Pahari, Bikash Kumar Behera, Manojita Dash, Richa Dash, Niranjana Sahoo, Indramani Nath, Aditya Prasad Acharya, Susen Kumar Panda, Subrata Mahapatra, Manoj V. Nair

## Abstract

The population of Asian elephants, an endangered species, is decreasing significantly due to anthropogenic interference in India. Research on Asian elephant disease is largely confined to observations reported from captive populations. The present study explored 11 disease investigation cases whose clinicopathological changes were similar to those of fatal hemorrhagic disease caused by *Bacillus anthracis*. We used molecular diagnostic approaches such as PCR (polymerase chain reaction), quantitative PCR, 16S metagenomics analysis and whole-metagenome sequencing for identification of the etiological agent responsible for these wild elephant mortalities. 16S metagenomics analysis revealed the presence of *Clostridium hemolyticum* as the dominant bacteria in all the mortality events. In contrast, flagellin gene-specific identification of all the samples and whole metagenomics analysis of blood samples collected from the ear pinna of a freshly deceased elephant revealed *Clostridium novyi* type B as the sole pathogenic agent responsible for the death of elephants. This study provides the first evidence of the involvement of *Clostridium novyi* type B in the death of wild elephants with similar clinical manifestations to anthrax. The current study also provides ample evidence of the need to explore investigations of emerging diseases of wild elephants to conserve these large mammals.

## 1. Introduction

The Asian elephant (*Elephas maximus*) is considered the only living species of the *Elephas* genus of the Elephantidae family. The Indian elephant (*Elephas maximus indicus*), which is a subspecies of Asian elephant, was designated an endangered species in 1976 by the International Union for Conservation of Nature (IUCN) on the Red List of Threatened Species [1]. On the basis of the data available for the year 2018, the global free-range Asian elephant population size is estimated to be approximately 48,323–51,680 individuals [2]. More importantly, India supports an estimated 60% of the global Asian elephant population and harbors nearly 29,964 Asian elephants spread over an area of 110,000 km2 of which 65,000 km2 is constituted as Elephant Reserves (ER). There are 33 ERs across 23 states, covering 11 elephant landscapes in four distinct regions, connected regionally by 150 corridors. The ERs cover approximately 30% of Protected Areas (PA), 40% of Reserved Forests (RF) and another 30% of private lands [2]. The anthropogenic intervention in natural habitats, fragmentation of forest corridors, electrocution, train accidents, human-elephant conflicts, infighting/falling, poisoning, and poaching have contributed to the significant decline in India’s free-range elephant population. However, mortality due to diseases are emerging as a significant threat to free-ranging elephants, especially fatal diseases such as anthrax [2] and elephant endotheliotropic herpesvirus hemorrhagic disease (EEHV-HD) [4, 5 & 6].

In addition to the above reported diseases, the current study identified another significant pathogen, *Clostridium novyi* type B, that can cause a ruinous effect on the health of the free-ranging elephant population. It is well documented that *C. novyi* type B causes infections in domestic animals, most notably infectious necrotic hepatitis (INH) in horses, sheep, pigs, and goats [7]. *C. novyi* type B is a strictly anaerobic bacterium that is present in the environment as a bacterial endospore and resistant to harsh abiotic factors. The endospore usually reaches mostly phagocytic liver cells as well as the spleen and bone marrow of herbivores through the portal circulation and remains dormant until a favorable anaerobic microenvironment develops [8]. The pathophysiology of INH is linked to initiating hepatic damage, usually induced by the migration of immature forms of liver flukes such as *Fascioloides* spp. and *Dicrocoelium* spp., which creates anaerobic conditions for the germination of spores and the proliferation of *C. novyi* type B [9]. No research has been conducted to explore the contribution of *C. novyi* type B as a pathological agent in large mammals, i.e., elephants. There are few studies available that discuss the involvement of *Clostridium perfringens* as the pathogen of captive Asian elephants [10, 11 & 12]. *C. perfringens* can cause fatal diseases along with coinfection with EEHV-4 in Asian elephants [12]. To understand the pathological etiology and identify previously undermined pathogens of the wild elephant population, advanced molecular tools are needed in wildlife disease investigations, especially emerging diseases.

The current study was designed to explore possible pathogen complexity in elephant disease investigations through a retrospective approach in wild elephants of Odisha, India.

## 2. Materials and methods

### 2.1. Sample collection

A total of 11 wild elephant death events were recorded during the study period (July 2021-- December 2022) and screened (Figure 1 & Supplementary Table 1). On the basis of changes in external features such as the oozing of blood from natural orifices such as the anus, mouth, and trunk, the field veterinarians considered elephant carcasses to be anthrax-suspected. In such situations, blood from the ear vein was collected through a sterile syringe following proper biosafety measures and transported in the cold chain to the Centre for Wildlife Health (CWH) for laboratory analysis within 6–12 hrs. of collection. The blood samples were processed for molecular detection of *Bacillus anthracis* within 6 hrs. after receiving samples at the CWH by following the method described below. The findings were reported back to the field veterinarians to conduct post mortem (PM) on the carcass if the case was negative for anthrax, following established guidelines issued by the Project Elephant, Ministry of Environment, Forest & Climate Change, Government of India [13]. The tissues from internal organs such as the spleen, heart, lungs, liver, kidney, and bone marrow, along with blood collected from the heart, were stored in a sterile container and transported to CWH in the cold chain within 12–14 hrs. of PM to identify the etiological agent responsible for elephant mortality. On the other hand, for non-anthrax suspected cases, PM was conducted within 3–4 h after the carcass was sighted, and the collected tissues were transported to CWH as mentioned above. These samples were stored at -20°C for long-term storage.

**Figure 1:**
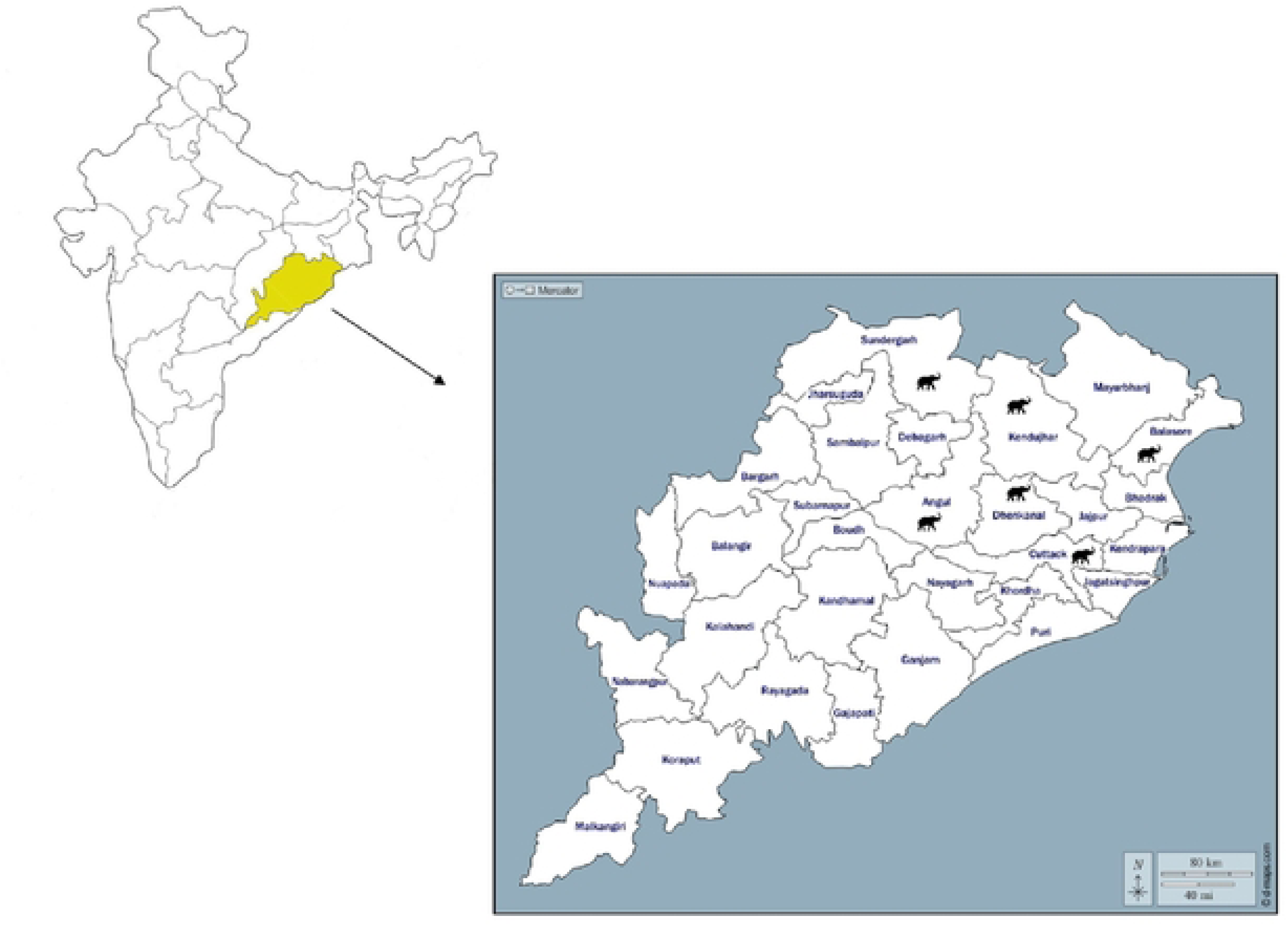
Geographical representation of elephant morality samples collected from different Forest Divisions of Odisha, India.

### 2.2. Extraction of DNA

The biological samples were processed in a biosafety cabinet for extraction of DNA. DNA extraction was performed via a PureLink Microbiome DNA Purification Kit (Invitrogen, Thermo Fisher Scientific, USA). The extracted DNA was quantified with a Qubit^TM^ fluorometer (Thermo Fisher Scientific, USA).

### 2.3. Molecular screening for *Bacillus anthracis*

The extracted DNA was used for real-time PCR for the detection of *B. anthracis* via specific primers that target the pXO1 (PA) and pXO2 (CAP) genes [14]. The amplification reactions were carried out in a total volume of 25 µl consisting of 5 µl (250 ng of DNA) of template DNA, 10 µM forward and reverse primers (**Table 1**), 2X real-time universal PCR master mix and molecular-grade nuclease-free water. The assay was performed in a Rotor-Gene Q instrument (Qiagen, Germany) with a protocol comprising 1 cycle at 95°C for 10 min followed by 40 cycles each at 95°C for 15 secs and 60°C for 1 min. Data analysis was performed via Rotor-Gene Q software, and Ct values were determined on the basis of the mean baseline signals during the early cycles of amplification. All reactions were run in triplicate, along with a negative control and synthetic DNA containing the PA and CAP genes as positive controls.

### 2.4. Molecular screening for elephant endotheliotropic herpesvirus type-1 (EEHV- 1)

All the samples were subjected to molecular screening for EEHV-1 to detect the presence of the herpes virus that causes viral hemorrhagic disease in elephants. The total DNA extracted from 11 disease samples through the previously mentioned method was used as the template for real-time PCR detection of EEHV-1 via specific primers targeting the EEHV-1 terminase gene [15]. The amplification reactions were carried out in a total volume of 25 µl consisting of 5 µl (250 ng of DNA) of template DNA, 10 µM specific primers (**Table 1**), 2X real-time universal PCR master mix and molecular-grade nuclease-free water. The assay was performed in a QuantStudio5 Real-Time PCR System (Applied Biosystems, USA) instrument with a protocol comprising 1 cycle at 500°C for 2 min and 95°C for 2 min followed by 40 cycles at 95°C for 15 sec and 60°C for 1 min. Synthetic DNA containing the EEHV-1 terminase gene sequence was used as a positive control. All reactions were run in triplicate, along with a negative control.

### 2.5. 16S rRNA gene metagenomics

Total DNA was extracted from different tissues via the method described above. The DNA was used as a template to amplify the *16S rRNA* gene. The V1-V9 variable regions of the *16S rRNA* gene were amplified via forward and reverse primers [16]. 16S rDNA PCR amplification was performed in a 25 μl volume using total DNA (10 ng), primers (5 nM), and Phusion High-Fidelity (HF) DNA Polymerase 5X master mix (Thermo Scientific, USA) according to the manufacturer’s protocol. The amplicons (∼1.6 kb) were analyzed on a 0.8% agarose gel. The PCR products were purified with Agencourt AMPure XP beads (Beckman Coulter, India) and eluted in 20 μl of buffer solution (10 mM Tris-HCl pH 8.0, with 50 mM NaCl). The quantities of the eluted products were determined via a Qubit 4 fluorometer (Thermo Scientific, USA) via a Qubit® 1X HS assay kit (Thermo Scientific, USA). A purified amplicon (100 fmol) was used as input DNA for multiplexed MinION-compatible library preparation via native barcodes (EXP-NBD104) and ligation sequencing (SQK-LSK109) following the manufacturer’s protocol (Oxford Nanopore Technologies, Oxford, UK). The pooled libraries were subsequently sequenced on a MinION Mk1C (ONT, Oxford, UK) using the R9.4.1 flow cell (MinION and Flongle flow cell; ONT, Oxford, UK), as per the manufacturer’s recommendation.

The final passed fastq sequences were uploaded to the EPI2ME 16S workflow online database with sequence sizes of 1000 to 2000 bp and a minimum identity of 80%. After completion of the analysis, the data were exported to a spreadsheet, and the results were compiled.

### 2.6. Identification of *Clostridium* species

After metagenomics analysis, a different approach was taken to identify the *Clostridium* species present in the tissue and blood samples collected from elephant mortality events. The extracted DNA was used as a template for multiplex conventional PCR, which identified the flagellin (*fliC*) genes of *Clostridium chauvoei*, *Clostridium hemolyticum*, *Clostridium novyi* types A and B, and *Clostridium septicum* [17] (**Table 1**). The PCR cycle was performed in a total reaction volume of 25 µl that contained multiplex 5X PCR master mix (NEB, USA), 5 nM of each primer, and 10 ng of template DNA. The amplified products were visualized via agarose (1.5%) gel electrophoresis. The amplified *fliC* gene was purified and Sanger sequenced, and the sequences were subsequently compared with those of other microorganisms via BLAST (http://blast.ncbi.nlm.nih.gov/Blast.cgi).

### 2.7. Whole-genome sequencing and analysis

The ear pinna blood sample collected from the CWH_540_Keonjhar case was selected for whole bacterial genome sequencing via the Illumina platform. Total DNA was extracted from a blood sample via a Gentra Puregene Blood Kit (Qiagen, Germany). The DNA was quantified via a Qubit fluorometer (Thermo Fisher Scientific, USA). A total of 50 ng of DNA was used to prepare the Illumina library via the Illumina® DNA Prepkit (Illumina, USA) following the manufacturer’s protocol. The final library was quality-checked through an Agilent tape station 4200 and sequenced on a NovaSeq 6000 (Illumina, USA) with 150 paired-end sequences.

The raw reads generated from NovaSeq 6000 sequencing were trimmed for adapter removal and low-quality reads (quality score <30 and length <20 bases) via the Trimgalore-v0.4.01 tool. The good-quality adapter-free reads were mapped against reference elephant mitochondrial and nuclear genomes via the Bwa-v0.7.5a2 tool to filter out host contamination. Approximately 30 million unmapped reads were retained after filtration. The unmapped reads were used for bacterial genome assembly in Unicyclerv 0.4.83. The assembled contigs were used for NCBI standalone blastv2.8.1+ 4 homology searches against 142 *Clostridium* genomes to filter out reliable contigs with the criteria of a minimum contig length of 200 bases, minimum identity, and coverage of 90% to generate the final assembled genome. The draft bacterial genome generated was used for prokaryotic gene/protein annotation via the prediction tool “Prokka-v1.145". The predicted proteins were annotated against the UniProt bacterial database with a minimum 30% identity cutoff for functional assignment and gene ontology through the DIAMOND BlastP6 program. Similarly, the predicted genes were also annotated against the KAAS7 database for pathway analysis with *Clostridium* species as a reference dataset. A DNA‒DNA hybridization analysis was also carried out for the assembled genome via the TYGS8 server. The average nucleotide identity of the assembled genome with the reference genome of *C. novyi* strain 150557 was calculated via the ANI Calculator9 [18]. The draft genome was used for the identification of AMR genes (Comprehensive Antibiotic Resistance Database, CARD) and virulence genes (Virulence Factor Database, VFDB). The mobile genetic elements present upstream and downstream of the AMR and VF genes were searched via the ACLAME database. The presence of secondary metabolite biosynthesis-related proteins was predicted through antiSMASH-v6.014 servers. The bacteriocin analysis was carried out between the predicted proteins of the assembled genome and the reference Bactibase database. Finally, the genetic variant annotation and functional effect prediction of the current bacterial genome were searched against the reference *Clostridium novyi* strain150557 through BCFtools and snpEFF.

### 2.8. Culture and Isolation

For further confirmation of the presence of *Bacillus anthracis,* the samples were processed for microbial culture examination [19 & 20]. The microbial culture of suspected *B. anthracis* samples was carried out in BSL2 facilities. The tissue samples from the suspected cases were inoculated on PLET agar (Polymyxin B - Lysozyme - EDTA – Thallous acetate Agar, manufactured by Sigma‒Aldrich, Germany) and sheep blood agar (Himedia) and incubated at 37°C for 48 h. The sample was considered positive for *B. anthracis* if at least a single colony was detected and identified by Gram-staining as well as 16S rRNA sequencing.

## 3. Results

The 11 elephant disease investigation cases included common gross pathological changes irrespective of sex and age. Briefly, these pathological changes included serosanguinous discharge from the eyes, bloody exudates from the trunk and other natural orifices, pale fascia, and petechial hemorrhagic musculature (Figure 2). During the postmortem examinations, the lungs, liver, kidney, and spleen were hemorrhagic and hyperemic in nature, and the heart was filled with unclotted blood, as well as petechial hemorrhages found on the pericardium, endocardium and ventricular septum. Postmortem examination revealed that the death of the elephant was due to septicemia.

**Figure 2:**
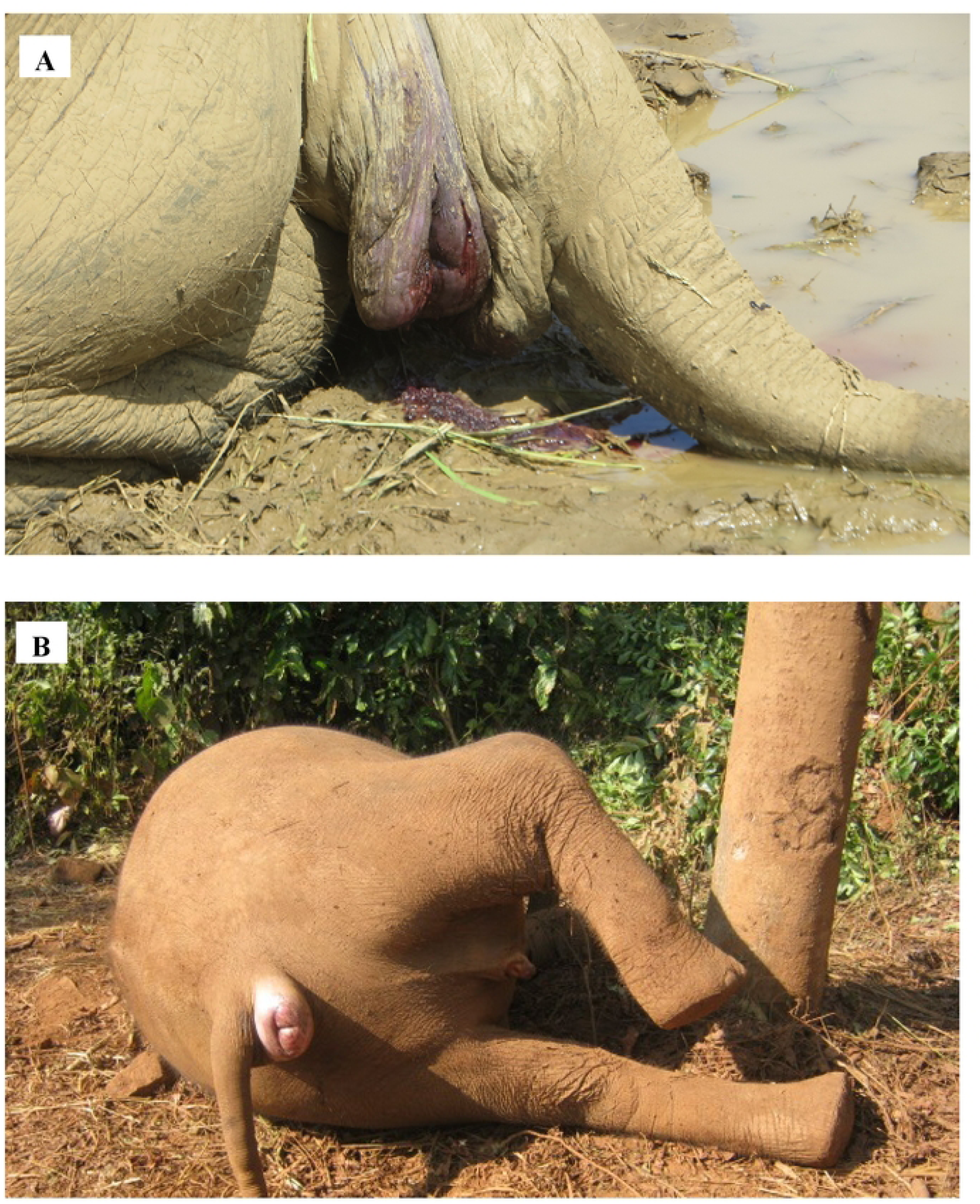
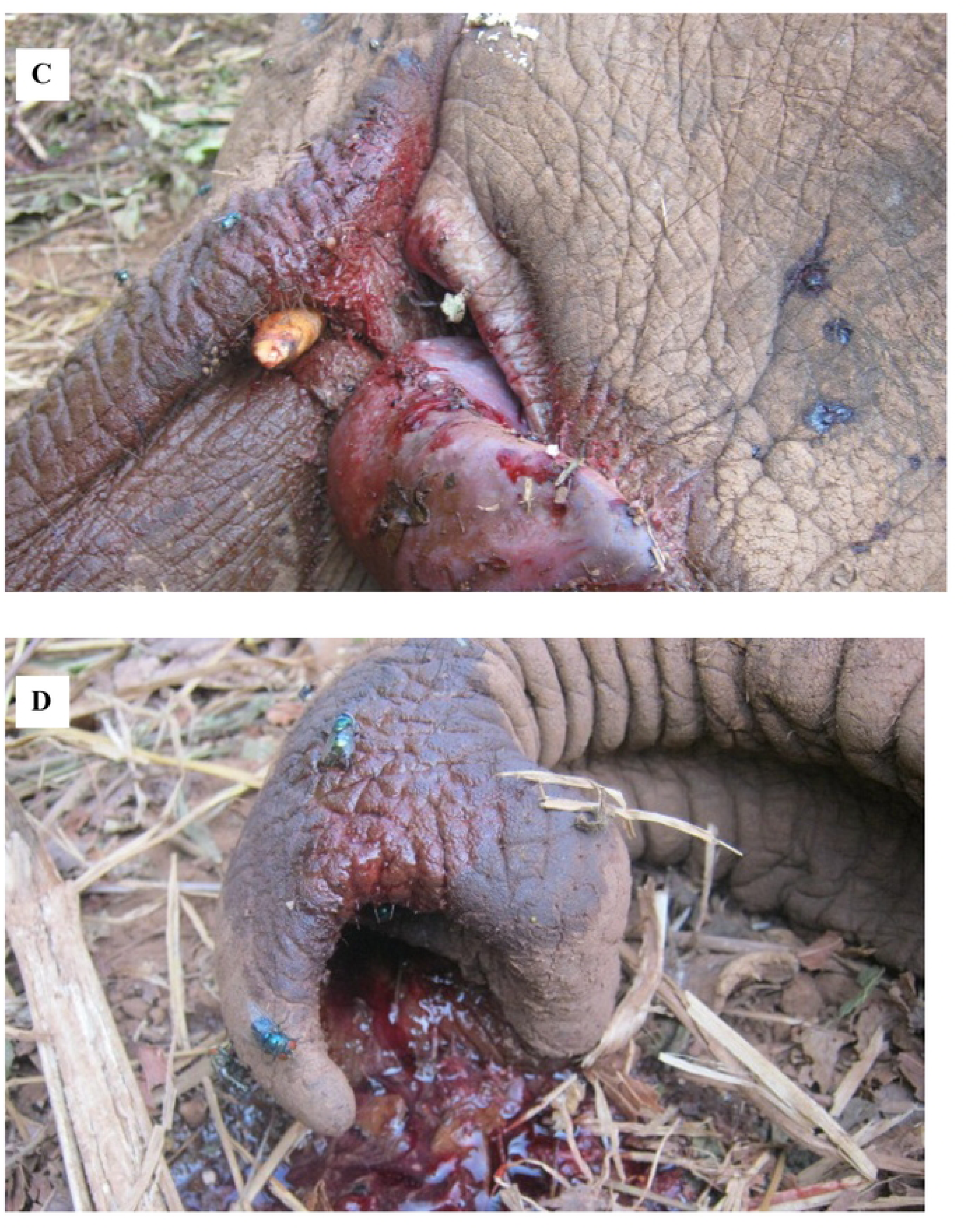
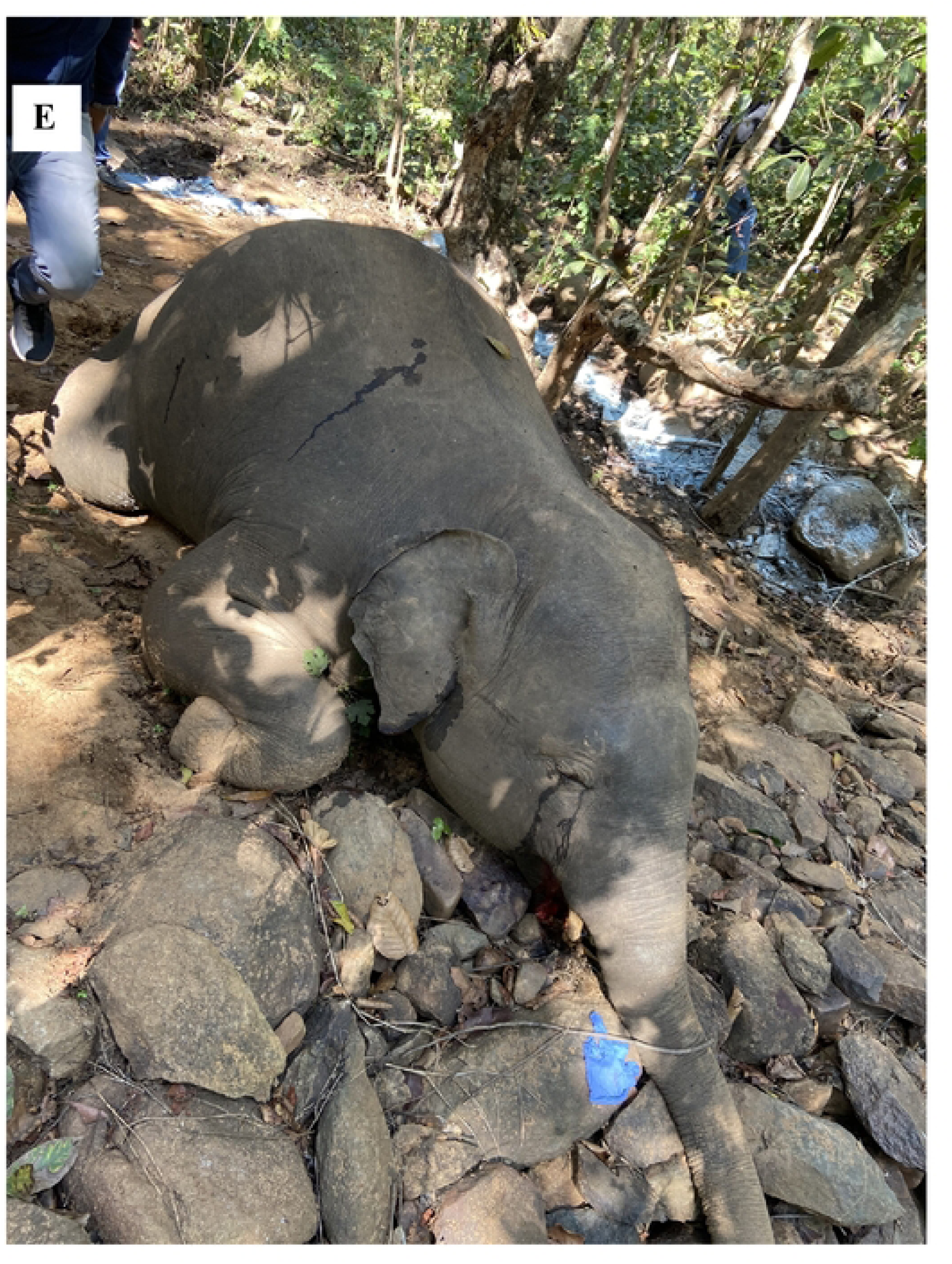
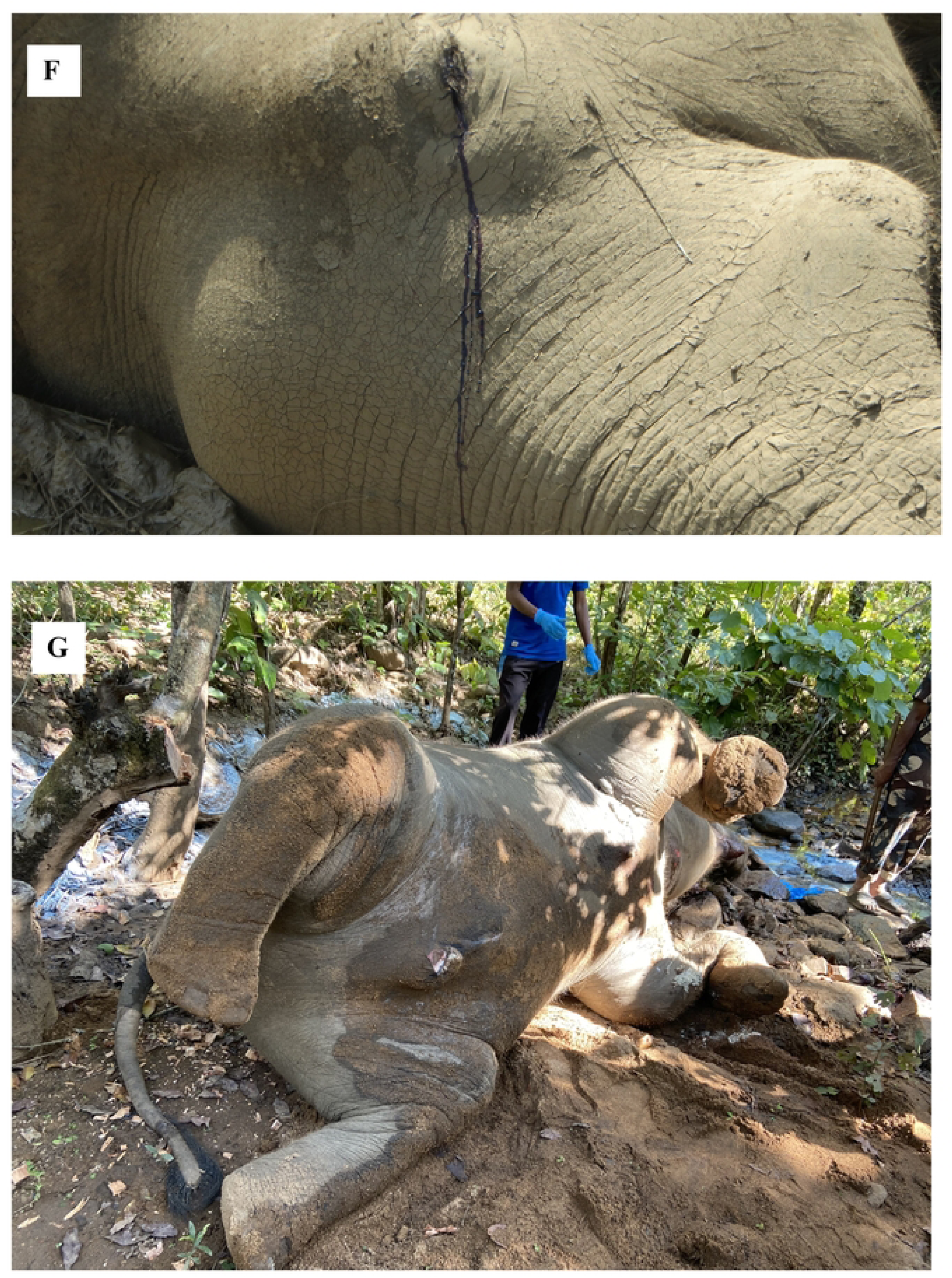
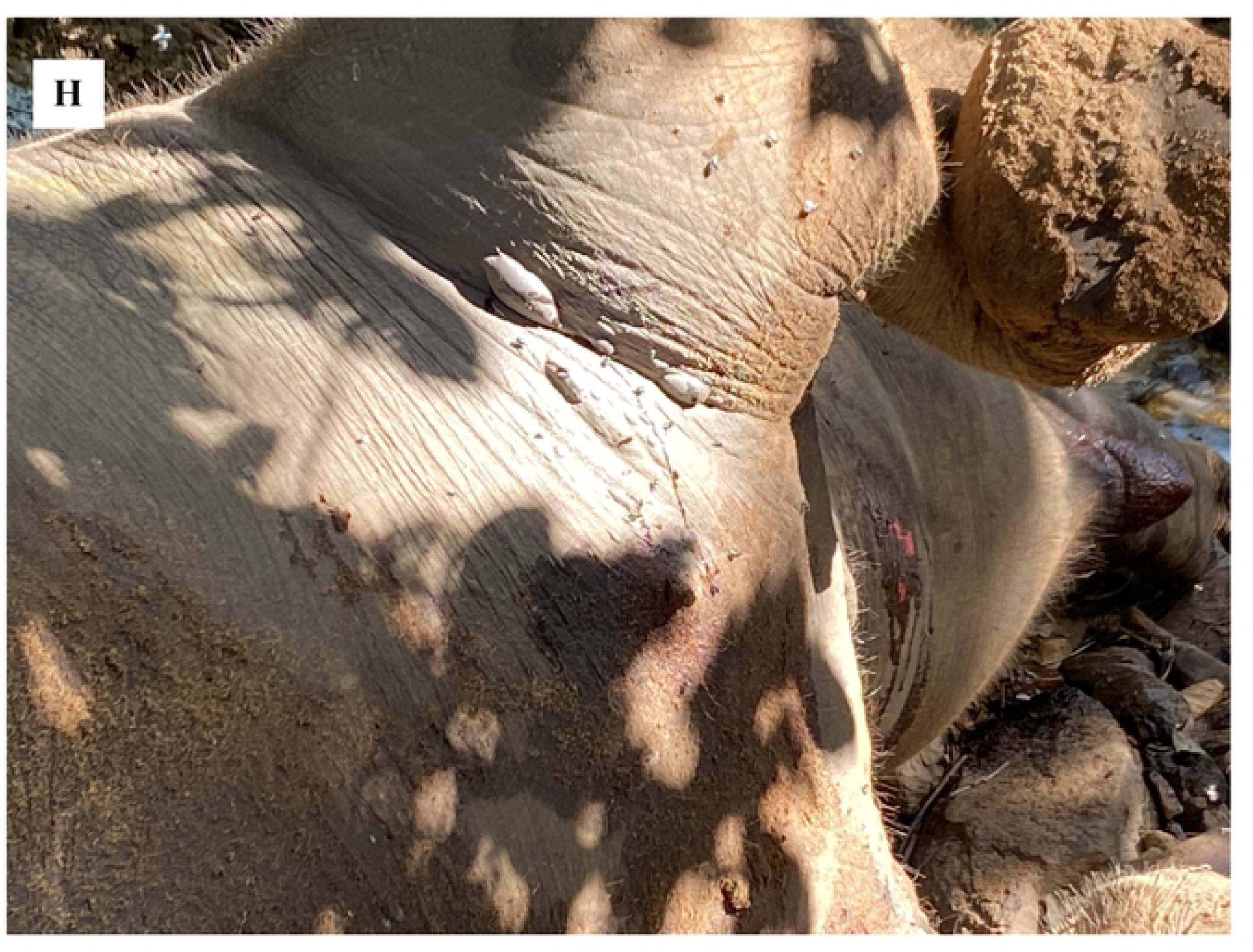
Gross morphological changes observed in the elephant carcasses. A- Bloody exudes from anus, B- Protrusion of anus, C & D – Un-clotted blood from mouth and trunk respectively, E & F- Serous discharged from the eye, G- Blotted carcass, picture taken after 12hr of death, H-Presence of gaseous gangrene at subcutaneous area of the forelimb

A total of 5 cases suspected for anthrax were screened against *Bacillus anthracis* and found to be negative through molecular assays as well as microbial culture. Similarly, EEHV 1 was negative for all the samples except for case CWH_540_Keonjhar (juvenile, 6–7 years old) (Supplementary Table 1).

A total of 11 individual elephant cases were processed for 16S metagenomics via the Oxford Nanopore Sequencing platform. A total of 117 to 13656 reads were generated for individual cases through sequencing (Supplementary Table 2). The generated reads were classified into *Clostridium hemolyticum*, *Clostridium* spp., Proteobacteria, and other bacterial species (Supplementary Table 2). The Proteobacteria group consisted primarily of *Burkholderia*, *Stenotrophomonas*, *Comamonas*, *Proteus*, *Shigella*, *Escherichia*, *Serratia, etc.,* and similarly, the other bacterial species groups included *Staphylococcus*, *Peptostreptococcus*, *Vagococcus, etc.* C*lostridium hemolyticum* (approximately 47.13% to 99.95% of the reads) was identified as the dominant bacterial population among the 10 disease investigation cases. In contrast to the above observations, the reads of the case “CWH_718_Dhenkanal” were classified into a diverse range of bacterial populations that were dominated by Proteobacteria (47.28%), along with *Clostridium hemolyticum* (32.01%). The raw reads were submitted under Bioproject PRJNA936996 with SRA accession numbers SRR23606898 to SRR23606906 and SRR23936849 to SRR23936851.

All the samples were amplified with a 427 bp product of the *fliC* gene via multiplex PCR. The PCR products were Sanger sequenced. A homology comparison of these sequences with those in the GenBank database revealed 99-100% similarity with the flagellin gene of *C. novyi* type B. The sequences were submitted to the NCBI database with accession numbers OQ628251-- OQ628261. A phylogenetic tree was constructed on the basis of the partial *fliC* sequences of 11 cases with other species of *Clostridium* (Figure 3).

**Fig 3:**
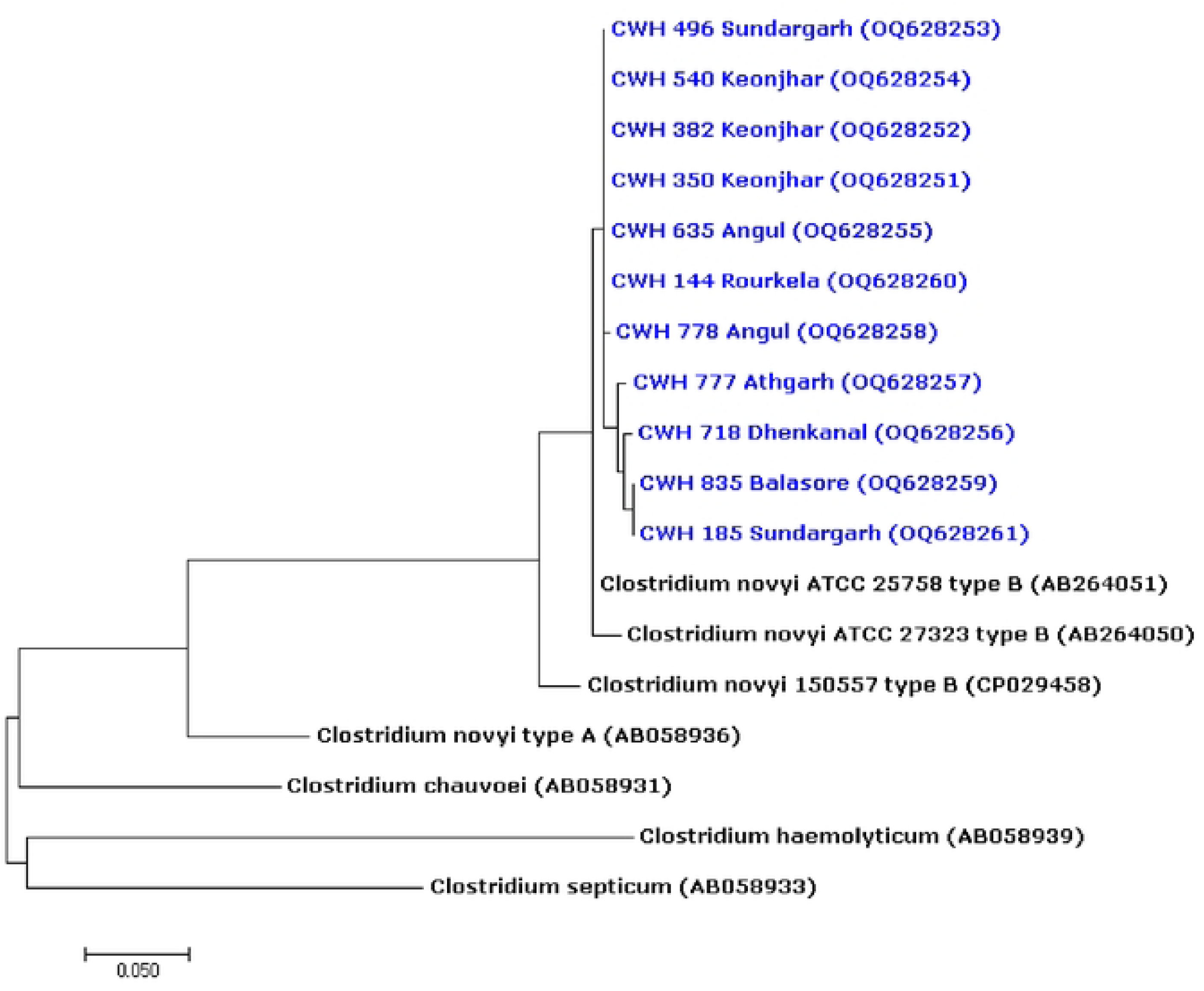
Phylogenetic tree based on comparison of partial flagellin gene (*fliC*) sequences using MEGA 7.0. The phylogenetic relationships between *Clostridium novyi* type B (Blue colored case ID with NCBI accession number) and other Clostridium species was generated using neighbour joining method with a Bootstrap values of 1000 replications.

The CWH_540_Keonjhar case tested negative for anthrax and showed the presence of *Clostridium hemolyticum* (79.06%) and *Clostridium* spp. (20.93%) in the 16S metagenomics analysis used for bacterial whole-genome sequencing. A total of 108 million Illumina paired-end short-read datasets were generated, and the sequencing data were submitted to the NCBI database under Bioproject Accession number PRJNA936629. Approximately 30 million unmapped reads were retained after host (elephant) read filtration. The unmapped reads were assembled and generated a total of 11.7 Mb of genome with 3,775 contigs. These assembled contigs were again filtered on the basis of the available 142 *Clostridium* genome homologs, which resulted in 3.1 MB of genome composed of 2873 contigs. Maximum homology was observed against the genome of the *C. novyi* strain 150557 (NCBI Bioproject Accession-PRJNA460203). A total of 2,424 proteins were predicted after the final draft genome annotation (Supplementary Figure 1).

DNA‒DNA hybridization analysis was used for the assignment of taxonomic status to the draft genome. The whole-genome phylogenetic results revealed that the draft genome of sample CWH_540_Keonjhar (7_A) was closely related to that of *C. novyi* strain 150557 (RefSeq assembly accession: GCF_003614235.1), with an average sequence identity of 90.8%. The average nucleotide identity was found to be 99.5% between the assembled genome of sample CWH_540_Keonjhar (7_A) and the reference genome of *C. novyi* strain 150557 via the ANI Calculator9 (Figure 4). The circular genome comparison of the assembled bacterial genome with that of the *C. novyi* strain 150557 showed similarity with the central reference sequence and other sequences as a set of concentric rings. BLAST matches are colored on a sliding scale indicating a defined percentage identity (Figure 5).

**Fig 4:**
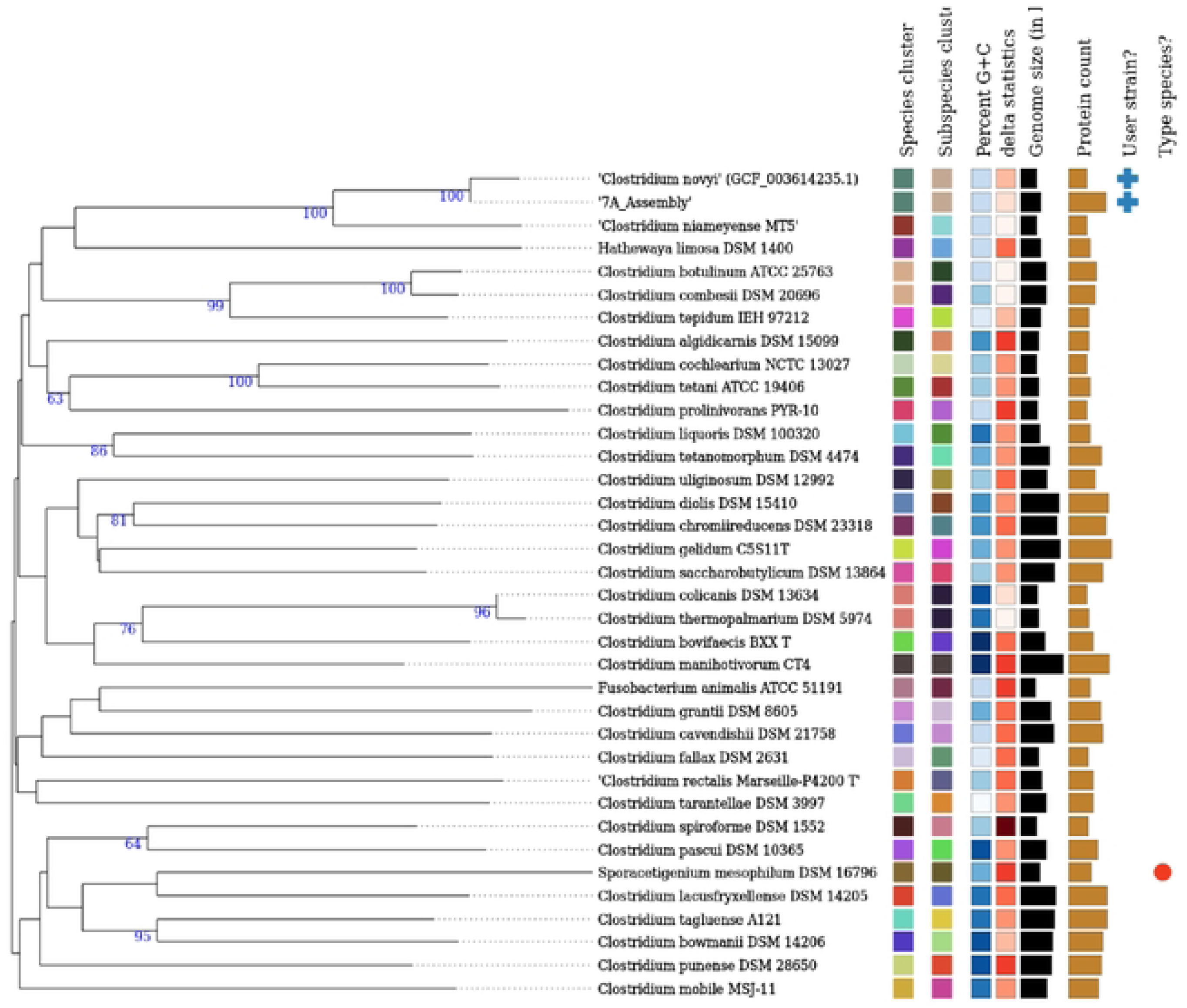
DNA-DNA hybridization analysis of draft assembly of CWH_540_Keonjhar (7_A) with *Clostridium novyi* strain 150557 (RefSeq assembly accession: GCF_003614235.1)

**Fig 5:**
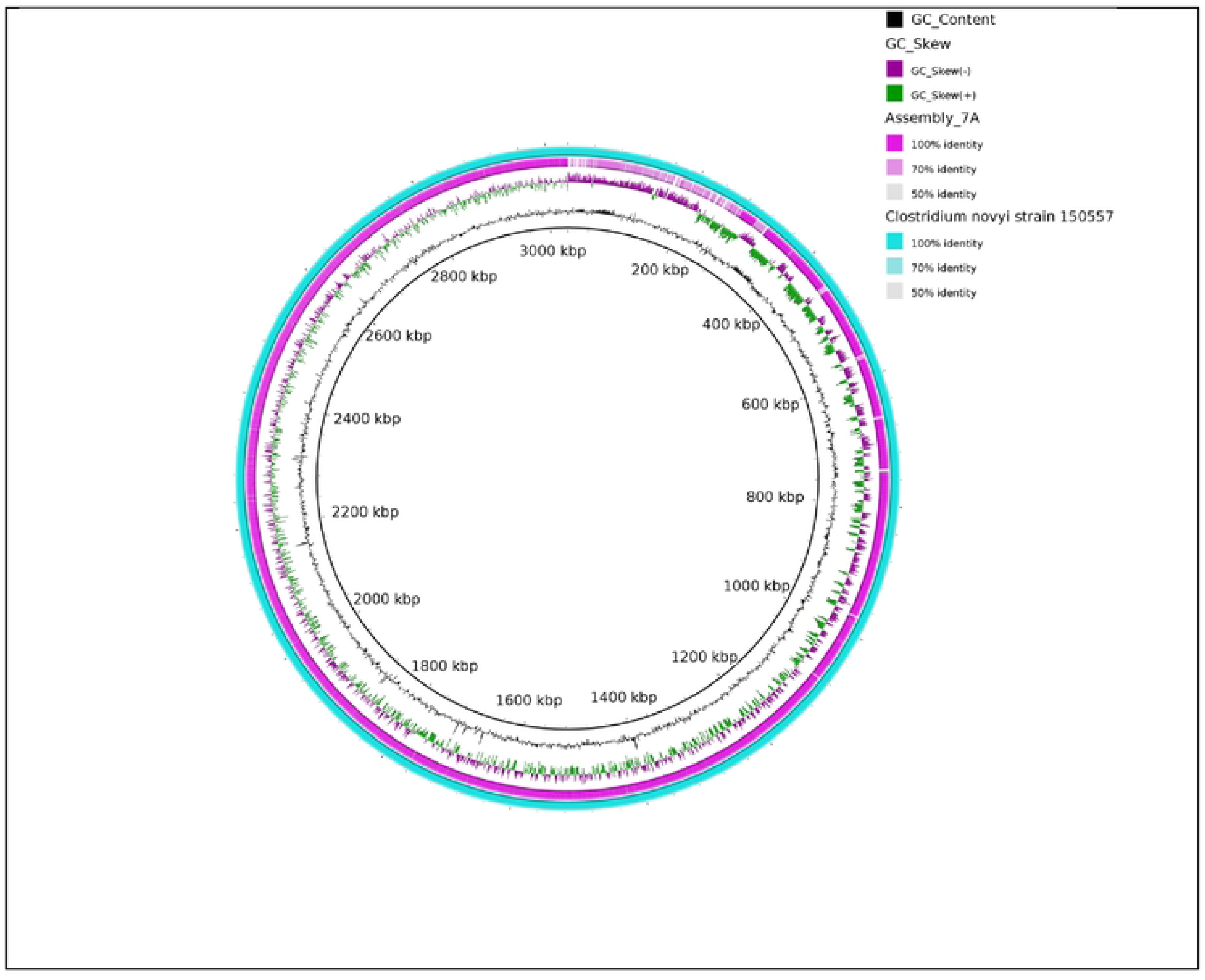
Circular genome comparison of the assembled bacterial genome i.e. CWH_540_Keonjhar (7_A) with *Clostridium novyi* strain 150557

Three significant antimicrobial-resistant protein-producing genes, rifamycin-resistant beta-subunit of RNA polymerase (*rpoB*), fluoroquinolone-resistant (*gyrB*), and elfamycin-resistant (*EF-Tu*), were identified in the draft genome. Similarly, a total of 7 genes responsible for the production of virulence factor proteins (alpha-clostripin precursor, fibronectin-binding protein, hemolysin III, chaperonin GroEL, tetanolysin O, elongation factor Tu and glyceraldehyde-3- phosphate dehydrogenase) were identified in the genome. The upstream and downstream regions (500 nucleotide sequence) of these AMR and VF proteins lacked mobile genetic elements (MGE). Genes encoding secondary metabolites and bacteriocin proteins were also absent in the genomes. Finally, the snpEFF tool was used for the prediction of 49,959 variants, of which 17,318 variants were reported to be nonsynonymous by annotation.

## 4. Discussion

India supports an estimated 60% of the global Asian elephant population and harbours nearly 29,964 elephants spread over an area of 110,000 km^2^ spread across four populations – North-western, Eastern, North-eastern and Southern. In India, the majority of free-range elephant mortalities occur due to poaching, electrocution, and sometimes due to diseases that are either EEHV or anthrax [21 & 22]. Therefore, considering the importance of preserving these endangered species also declared as India’s National Heritage Animal, apart from other measures, a spatial disease investigation study that provides insight into the health status of the remaining elephant population of India is also highly necessary.

Subadult elephants, i.e., those younger than 10 years, are more susceptible to fatal hemorrhagic diseases caused by EEHV [23]. However, in the present study, only one of the 3 subadult elephant cases whose ages were younger than 10 years was positive for EEHV1 infection. Apart from that in India, intense competition for resources is present between livestock and free-range elephants near areas prone to anthrax [3]. Although all the anthrax suspected elephant carcasses found in the geographical anthrax suitability area in the present study [3], none of them were positive for *Bacillus anthracis*.

A total of 11 elephant cases were screened via 16S metagenomics analysis to detect potential bacterial pathogens involved in animal mortality. 16S metagenomic sequencing revealed the presence of *Clostridium hemolyticum* as the dominant bacteria in all the mortality cases, with various numbers of reads. These findings indicate that *Clostridium hemolyticum* might be the primary pathogen responsible for elephant death. In contrast to the above observations, whole-genome sequencing was conducted on one of the samples to obtain complete genomic profiling of the suspected pathogen. *Clostridium novyi* type B was detected through whole-genome sequencing. The contradiction between the 16S metagenomics and whole-genome results can be explained by the fact that *Clostridium hemolyticum* and *Clostridium novyi* type B have very similar biological characteristics. The molecular identification of these bacteria on the basis of 16S rDNA sequencing is often considered difficult, as the sequences are completely identical [17]. Additionally, flagellin gene typing validated the presence of *C. novyi* type B in all the deceased animals. The culture and isolation of pathogenic *Clostridium* spp. are considered difficult, as bacteria require strict anaerobic conditions to grow, and the biological samples collected postmortem are often contaminated with other anaerobic bacteria, including other *Clostridia* originating from the intestine and soil [17]. Interestingly, during 16S rDNA sequencing, out of the 11 elephant mortality cases, 7 presented *Proteobacteria* and other bacterial species apart from the *Clostridia* group. This observation signifies that the decomposition of carcasses had already started before necropsy. In general, accelerated body decomposition is often observed in animals that are associated with *Clostridial* diseases. Notably, for the correct diagnosis of *Clostridium* infection, rapid evaluation and preservation of tissues after the death of the animal becomes mandatory [24]. In the present study, swift collection of fresh tissue samples from recently deceased free-range elephants was a challenging task, as most of the time, rigorous surveillance of ill animals could not be achieved in dense forest regions.

According to postmortem studies on deceased elephants, the internal organs, including the lungs, liver, heart, spleen, and kidneys, are either engorged or hemorrhagic. Importantly, *C. novyi type* B infection can result in liver bleeding, the formation of fibrin and cell debris, and focal to multifocal coagulative necrosis [24]. Interestingly, in the present study, the necropsy reports of dead elephants from field veterinarians yielded comparable findings, i.e., vascular congestion, multifocal parenchymal and capsular hemorrhages, and fibrin thrombi in the liver and heart blood vessels, as described by Nyaoke et al. [24] in horses.

*C. novyi* type B is frequently referred to as a hardy bacterium because of the extreme environmental resistance of the spores. The spores are regularly found in soil and still bodies of water, and there is a significant likelihood that grazing animals will consume them regularly [25 & 26]. After entering the digestive system, *C. novyi* type B spores are absorbed from the intestine and transferred to the liver via the portal circulation before moving on to other organs. Additionally, spores are phagocytosed and persist in a dormant state inside bone marrow macrophages, spleen Kupffer cells, and the liver [25 & 27]. Most of the elephant’s internal organs (liver, spleen, kidney, and bone marrow) demonstrated the active presence of *C. novyi* type B, which further supported the congruency of the observations described above. In addition, the field veterinarians in this investigation observed serosanguinous oozes from the natural orifices of dead elephants, such as anthrax-suspected mortalities. This observation focused on the fact that the vegetative forms of *C. novyi* type B secrete toxins that significantly alter and disrupt the actin cytoskeleton as well as the vimentin and tubulin system of the capillary endothelium [28 & 29], causing fluid to extravasate in the peritoneum and from natural orifices [30].

According to the general consensus, infectious necrotic hepatitis develops following liver damage, resulting in necrosis and the accompanying anaerobic conditions necessary for the germination of dormant *C. novyi* type B spores and the synthesis of toxins [30]. The most significant risk factor for INH in sheep, cattle, goats, buffalo, camelids, and cervids is thought to be the migration of immature liver flukes through the liver. In the present study, the involvement of liver flukes in creating a hepatic anaerobic microenvironment in deceased elephants was not reported by field veterinarians during necropsy. However, some studies have shown that *Fasciola jacksoni* is present in the bile ducts of free-range Asian elephants during postmortem examinations [31 & 32]. The occasional occurrence of INH in animals without liver flukes has been documented in a few cases. Other potential causes of liver injury have been proposed in those circumstances, although their contribution to the predisposition to INH has not yet been clearly established [8].

## 5. Conclusion

In conclusion, a thorough investigation is essential to understand the pathogenic modalities of *C. novyi* type B since the organism is difficult to identify and there is a shortage of knowledge on *C. novyi* type B infection in wild animals such as elephants. Owing to the genetic similarity between *C. novyi*, *C. botulinum*, and *C. hemolyticum*, the organism was incorrectly identified by 16S metagenomic sequencing and could not be conclusively identified. Subsequent tests, such as whole-genome sequencing, were able to identify *C. novyi* type B more definitively as the etiologic agent in current elephant motility cases. Furthermore, the strict anaerobic nature of *C. novyi* makes it challenging to grow in the laboratory and extract high-quality DNA for whole-genome sequencing. This incident also emphasizes the value of advanced molecular diagnosis, which provides discrimination between *Clostridial* species. The current research highlights the need for an advanced disease investigation strategy that is needed to conserve threatened wild animals throughout the world. The risk posed by pathogens can potentially impact not only elephant populations but also ecosystems as a whole. Moreover, understanding the cause of death is crucial for implementing appropriate preventive and control measures to protect elephant populations. Additionally, a synchronized collaboration between government agencies and the public through awareness creation and education about the importance of wildlife conservation is highly important.

## Acknowledgment

The research work was funded by the Forest, Environment and Climate Change Department, Government of Odisha, India.

## Ethical statement

This study was conducted on dead wild elephants, found in arcane forest. The post mortems were conducted by veterinarian by following established guidelines issued by the Project Elephant, Ministry of Environment, Forest & Climate Change, Government of India. No institutional ethical committee was necessary for this study.

## Author Contributions

S.K. Padhi, N.S., I.N., A.P.A and S.K. Panda designed the research; S.K. Padhi, A.P., B.K.B., M.D., and R.D. performed the research; S.K. Padhi, N.S., S.M., I.N. and S.K. Panda contributed new reagents/analytic tools and sample collection; S.K. Padhi analyzed the data; and S.K.Padhi, A.P., N.S. and M.V.N. wrote the paper.

## Supporting information

**Supplementary Table 1.xlsx:** Detailed summary of dead elephants

**Supplementary Table 2.docx:** 16s metagenomics analysis data generated through Oxford Nanopore Technology (ONT) sequencing

